# Corneal Biomechanical Alterations in Patients with Chronic Ocular Graft Versus-host Disease

**DOI:** 10.1101/553867

**Authors:** Giuseppe Giannaccare, Marco Pellegrini, Leonardo Taroni, Federico Bernabei, Carlotta Senni, Arianna Grendele, Vincenzo Scorcia, Emilio C Campos

## Abstract

**Purpose:** To compare corneal biomechanics between patients with ocular graft versus-host disease (oGVHD) and healthy subjects (controls), and to further correlate these values with ocular and hematological characteristics.

**Materials and Methods:** The following procedures were performed in oGVHD patients and controls: Schirmer test (ST), break-up time (BUT), corneal and conjunctival staining, tear matrix metalloproteinase-9 (MMP-9) assay (InflammaDry test, Rapid Pathogen Screening, Inc, Sarasota, FL). Corneal biomechanics were calculated by using ocular response analyzer (ORA, Reichert Instruments, Depew, New York, USA). The Mann-Whitney U test was used to compare continuous variables between oGVHD patients and controls. Correlations of corneal biomechanics with ocular and hematological parameters were examined using Spearman’s correlation.

**Results:** A total of 45 oGVHD patients (mean age ± SD, 51.5 ± 7.1 years) and 34 controls (47.8 ± 6.1 years) were included. Patients with oGVHD showed significantly lower values of corneal hysteresis (CH) and corneal resistance factor (CRF) compared to controls (respectively, 9.4 ± 1.8 mmHg vs 11.6 ± 1.6 and 9.7 ± 1.4 mmHg vs 12.3 ± 1.3; always *p<*0.001). Twenty-nine of the oGVHD eyes (64.4%) were strong-positive for MMP-9, while 16 (35.6%) were weak-positive. Conversely, only 4 of the control eyes (11.8%) were weak-positive for MMP-9. In patients with oGVHD, CH was significantly correlated with corneal staining (*Rs* = −0.316, *p* = 0.035), conjunctival staining (*Rs* = −0.437, *p* = 0.003), ST (*Rs* = 0.390, *p* = 0.008), BUT (*Rs* = 0.423, *p* = 0.004), oGVHD severity grade (Rs = −0.383, *p* = 0.009), and MMP-9 positivity grade (*Rs* = −0.429, *p* = 0.003), while CRF was correlated only with corneal staining (*Rs* = −0.317, *p* = 0.034).

**Conclusions:** Corneal biomechanics are reduced in patients with oGVHD, and CH is negatively correlated with disease severity grade and MMP-9 tear levels.

## Introduction

Allogeneic hematopoietic stem cell transplantation (HSCT) is an established and potentially curative treatment for a variety of malignant and non-malignant hematological disorders. Graft versus-host disease (GVHD) is a multi-organ systemic disease caused by complex interactions between donor and recipient immune systems and represents the leading cause of morbidity following HSCT.[1] Chronic ocular GVHD (oGVHD) develops in 40 to 60% of patients undergoing HSCT, and dry eye disease (DED) represents the hallmark of this condition.[2–4] The disease is thought to be the result of the progressive immune-mediate inflammatory damage of ocular surface structures, which may lead to lacrimal and meibomian glands dysfunction, conjunctival keratinization, corneal epitheliopathy, eyelid laxity and scarring, and in more severe cases even corneal melting and perforation.[5–7]

Recent proteomic studies evaluated the various constituent compositions of tears inflammatory mediators in eyes with oGVHD and detected increased levels of cytokines and matrix metalloproteinases (MMPs).[8,9] Among these, MMP-9 is a proteolytic enzyme involved in the degradation of corneal extracellular matrix components, whose dysregulation is implicated in impaired wound healing and stromal keratolysis.[10–12] The excessive proteolytic activity in the cornea determines the degradation of stromal components, that are known to be the main determinants of biomechanical corneal behavior. Thus, we hypothesized that higher tears levels of MMP-9 might affect the ultrastructure of the corneal stroma, producing changes in corneal biomechanics.

In the present study, we compared the corneal biomechanical properties in patients with oGVHD and healthy matched subjects; in addition, we investigated whether these properties were correlated with ocular surface and hematological characteristics of oGVHD patients.

## Materials and Methods

### Study Design and Patients

This prospective case-control study was conducted at a single tertiary-referral Ocular Surface Center (S.Orsola-Malpighi University Hospital, Bologna, Italy) between January 2018 and October 2018. The study was performed in accordance with the principles of the Declaration of Helsinki and was approved by the local Institutional Review Board. Written informed consent was obtained from all subjects included in the study before any procedure.

Consecutive hematological patients who underwent allogeneic HSCT and came at our Center for routinely ocular surface check-up visits were screened for enrollment. Inclusion criteria were: age older than 18 years and a diagnosis of chronic oGVHD according to the International chronic ocular GVHD Consensus Group.[13] The severity of oGVHD was graded as none (grade 0), mild/moderate (grade I) and severe (grade II).[13] Healthy sex- and age-matched subjects acted as controls. All the study procedures were carried out in one eye for both patients and controls according to an automatic randomization procedure (www.randomization.com).

Exclusion criteria for both groups included any prior ophthalmologic surgery in the last year, keratoconus, corneal dystrophy, contact lens wearing, glaucoma, spherical equivalent ≥ 5 diopters, astigmatism ≥ 3 diopters and any ocular infection within 3 months prior to enrollment.

Clinical data including demographic characteristics, ongoing systemic and topical therapies, hematological diagnosis, duration of disease, type of HSCT donor, time from GVHD diagnosis to ophthalmological evaluation, presence and localization of systemic GVHD, and history of previous therapies (i.e. autologous HSCT, radiotherapy [RT], and chemotherapy [CT]) were collected at the time of the ophthalmic evaluation.

### InflammaDry Testing

The InflammaDry test (Rapid Pathogen Screening, Inc, Sarasota, FL) was performed according to the manufacturer’s instructions to determine whether both oGVHD patients and control subjects exhibited pathological levels of MMP-9.[14] The use of any topical ophthalmic medications was discontinued two hours prior to examination. In brief, a tear sample was collected by lowering the palpebral conjunctiva and gently dabbing the fleece of the sample collector 8 to 10 times in multiple locations, allowing the patient to blink between dabs to ensure saturation. The sampling fleece glistened or turned pink when an adequate sample was collected, and was then snapped into the test cassette. The absorbent tip was immersed into the buffer vial for 20 seconds and laid flat on a horizontal surface for 10 minutes before interpretation of test results. The presence of one blue line and one red line in the test result window indicated a positive test result (MMP-9 ≥ 40 ng/mL), whereas only one blue line indicated a negative test result (MMP-9 < 40 ng/mL). In addition, the positive red line was compared with the grading index to classify the result as weak-positive and strong-positive.[15] Three well-trained observers (FB, CS & AG) individually interpreted the InflammaDry grading result, masked to both subject characteristics and clinical diagnosis. The result was confirmed when the interpretations of at least two of the observers were in agreement.

### Ocular Surface Work-up

Subjective symptoms of ocular discomfort were scored by the Ocular Surface Disease Index (OSDI) questionnaire. Slit-lamp examination with ocular surface staining was performed after administration of 2mL of 2% fluorescein dye using the blue cobalt filter and a 7503 Boston yellow filter kit to enhance staining details. Corneal and conjunctival staining were graded using respectively the Oxford score[16], and the van Bijsterveld score[17]; Schirmer test type I and break up time (BUT) measurements were performed according to the Dry Eye Workshop guidelines.[18]

### Ocular Response Analyzer Measurements

The ocular response analyzer (ORA, Reichert Instruments, Depew, New York, USA) measures two applanation pressure points during a dynamic bi-directional applanation process generated by a precisely metered air pulse. The first pressure point occurs as an air puff pushes the cornea inward, while the second pressure point as the cornea returns from the applanated state to normal. The difference between these two pressures is defined as the CH, while CRF is calculated as a linear function of the two values. The average of both pressures provides Goldmann-correlated IOP (IOPg), while the corneal-compensated IOP (IOPcc) is the recalculated IOP value using corneal biomechanical information provided by the CH measurement.[19]

All the ORA examinations were performed by the same operator (LT) blinded to subject’s characteristics. The use of any topical ophthalmic medications was discontinued two hours prior to examination. Before each examination, central corneal thickness (CCT) values obtained with an ultrasonic pachymeter (Dicon P55, Paradigm Medical Industries Inc., Salt Lake City, UT, USA) were inserted in the software. Because of the potential confounding effect of diurnal IOP variation, all measurements were obtained between 10 AM and 12 PM. All included ORA measurements had a waveform score > 7.0.[20] The average values of four measurements with desirable curves were recorded for statistical analysis.

### Statistical Analysis

The SPSS statistical software (SPSS Inc, Chicago, Illinois, USA) was used for data analysis. Values are expressed as mean ± standard deviation (SD). The Mann-Whitney U test was used to compare continuous variables between ocular GVHD patients and control subjects. The χ^2^ test was used to compare dichotomous variables between ocular GVHD patients and control subjects. Correlations of corneal biomechanical properties with demographical, hematological and ocular parameters in the oGVHD group were examined using Spearman’s correlation analysis. A Bonferroni correction for multiple testing was used by multiplying the observed *p*-value with the number of comparisons within each analysis. A *p* value < 0.05 was considered statistically significant.

## Results

We screened a total of 51 post-HSCT patients during the study period. Of these, 45 patients fulfilled the inclusion criteria, and were finally enrolled in the study: 5 of them (11.1% of the total) belonged to the oGVHD severity grade 0, 11 (24.4%) to grade I and 29 (64.4%) to grade II. The remaining 6 patients were excluded from the analysis because the diagnosis of oGVHD was not reached (n = 3), or due to the presence of significant corneal alterations (neovessels) hampering accurate ORA measurements (n = 3). Thirty-four healthy subjects were enrolled as control group. The demographic and clinical parameters of patients included in the analysis are reported in Table 1. No significant differences in age and sex distribution between hematological patients and control subjects were found (always *p* > 0.05).

**Table 1:** Demographic and clinical parameters of patients with ocular GVHD.

| Characteristic | Number (%) |
| --- | --- |
| Patients | 45 |
| Males | 22 (48.9) |
| Females | 23 (51.1) |
| Age (years) | 51.2 ± 7.1 |
| Hematological diagnosis |  |
| AML | 14 (31.1) |
| ALL | 12 (26.7) |
| HL | 2 (4.4) |
| Non-HL | 4 (8.8) |
| CML | 9 (20) |
| CLL | 2 (4.4) |
| MM | 2 (4.4) |
| Systemic GVHD |  |
| None | 4 (8.8) |
| Skin | 28 (62.2) |
| Lung | 14 (31.1) |
| Mouth | 17 (37.8) |
| G-I tract | 12 (26.7) |
| Genitalia | 5 (11.1) |
| Other | 9 (20) |
| Time from GVHD diagnosis to visit (months) | 65.9 ± 54.9 |
| Donor |  |
| Unrelated | 27 (60.0) |
| Related | 18 (40.0) |
| Previous hematological therapies |  |
| HSCT | 9 (20) |
| RT | 29 (64.4) |
| CT | 6 (13.3) |
| Ongoing systemic therapy |  |
| None | 14 (31.1) |
| Corticosteroids | 21 (46.7) |
| Other Immunosuppressants | 7 (15.5) |
| Antivirals/Antibiotics | 23 (51.1) |
| Other | 6 (13.3) |
| Ongoing topical therapy |  |
| Corticosteroids | 19 (42.2) |
| Cyclosporine | 6 (13.3) |
| Tear substitutes | 45 (100) |
| Blood-derived eye drops | 9 (20) |
| Other | 10 (22.2) |

GVHD: Graft versus-host disease; AML: Acute myeloid leukemia; ALL: Acute lymphoblastic leukemia; HL: Hodgkin lymphoma; CML: Chronic myeloid leukemia; CLL: Chronic lymphocytic leukemia; MM: Multiple myeloma; G-I: gastro-intestinal; HSCT: hematopoietic stem cell transplantation; RT: radiotherapy; CT: chemotherapy.

Ocular surface parameters of oGVHD patients and control subjects are reported in Table 2. A significant difference of all parameters was found between the two groups (always *p* < 0.05).

**Table 2:** Ocular surface parameters in patients with ocular GVHD and control subjects.

| Parameter | Ocular GVHD | Control group | *P* |
| --- | --- | --- | --- |
| OSDI score | 58.7 ± 21.4 | 7.1 ± 3.1 | < **0.001** |
| Schirmer test I (mm/5') | 2.9 ± 3.6 | 23.2 ± 8.6 | < **0.001** |
| BUT (s) | 1.9 ± 2.0 | 10.9 ± 3.1 | < **0.001** |
| Corneal staining (Oxford score) | 2.4 ± 1.4 | 0.2 ± 0.4 | < **0.001** |
| Conjunctival staining (VB score) | 9.4 ± 5.6 | 0.7 ± 0.5 | < **0.001** |

GVHD: Graft versus-host disease; OSDI: Ocular surface disease index; BUT: break-up time; VB: van Bijsterveld.

Patients with oGVHD showed significantly lower values of CH and CRF compared to controls (respectively, 9.4 ± 1.8 mmHg vs 11.6 ± 1.6 and 9.7 ± 1.4 mmHg vs 12.3 ± 1.3; always *p <* 0.001) (Table 3). In addition, CCT was significantly lower in oGVHD patients compared to controls (510.7 ± 32.5 μm vs 537.9 ± 29.8, *p* < 0.001).

**Table 3:** Corneal biomechanical properties and central corneal thickness in patients with ocular GVHD and control subjects.

| Parameter | Ocular GVHD | Control group | *P* |
| --- | --- | --- | --- |
| CH (mmHg) | $9.4 \pm 1.8$ | $11.6 \pm 1.6$ | **<0.001** |
| CRF (mmHg) | $9.7 \pm 1.4$ | $12.3 \pm 1.3$ | **<0.001** |
| IOPg (mmHg) | $16.2 \pm 5.0$ | $18.1 \pm 3.6$ | 0.054 |
| IOPcc (mmHg) | $17.7 \pm 5.6$ | $16.9 \pm 4.1$ | 0.527 |
| CCT ($\mu\text{m}$) | $510.7 \pm 32.5$ | $537.9 \pm 29.8$ | **<0.001** |

CCT: Central corneal thickness; GVHD: Graft versus-host disease; CH, corneal hysteresis; CRF, corneal resistance factor; IOPg; Goldmann-correlated intraocular pressure; IOPcc, corneal-compensated intraocular pressure.

The proportion of MMP-9 tear film positivity significantly differs between the two groups (*P* < 0.001). In particular, 29 of the oGVHD eyes (64.4% of the total) were strong-positive for MMP-9, while 16 (35.6%) were weak-positive. Conversely, only 4 of the control eyes (11.8%) were weak-positive for MMP-9 (Figure 1).

**Figure 1:**
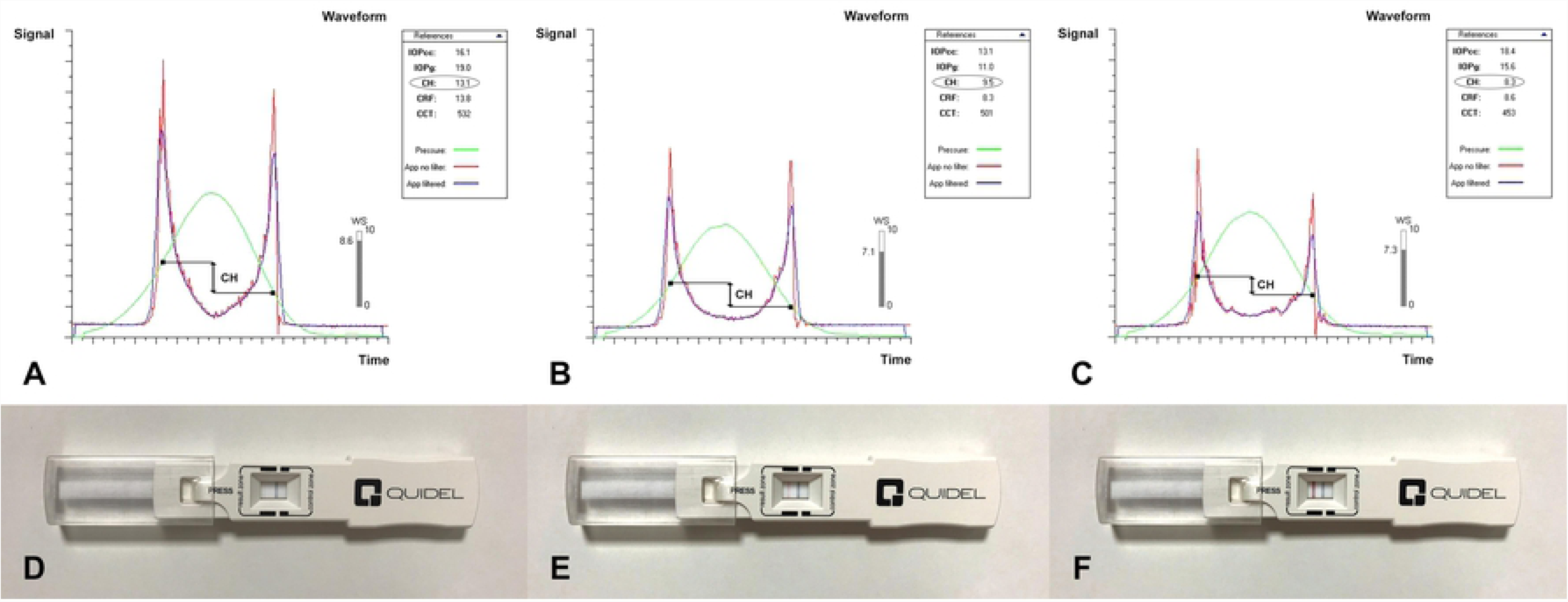
Ocular response analyzer measurement signals and InflammaDry assays of 3 representative subjects: healthy control, negative for MMP-9 (Parts A, E); ocular graft versus-host disease (oGVHD) patient (severity grade I), weak-positive for MMP-9 (Parts B, F); oGVHD patient (severity grade II), strong-positive for MMP-9 (Parts C, F).

In patients with oGVHD, CH was significantly correlated with corneal staining (*Rs* = −0.316, *p* = 0.035), conjunctival staining (*Rs* = −0.437, *p* = 0.003), Schirmer test (*Rs* = 0.390, *p* = 0.008), BUT (*Rs* = 0.423, *p* = 0.004), oGVHD severity grade (Rs = −0.383, *p* = 0.009), and MMP-9 positivity grade (*Rs*=-0.429, *p*=0.003), while CRF was correlated only with corneal staining (*Rs*=-0.317, *p*=0.034).

No significant correlations were found between CH and CRF values and hematological characteristics (always *p*>0.05).

## Discussion

The cornea is a complex biomechanical composite, and the collagen present in the Bowman’s layer and stroma provides the major contribution to its biomechanical behavior.[21,22] Frequently, oGVHD determines the damage of corneal epithelium that can be observed at slit lamp examination as punctate keratopathy, filamentary keratitis, and persistent epithelial defect. In more severe cases, the persistent impairment of wound healing alters the stromal ultrastructure, with the onset of keratolysis, corneal melting and even perforation.[23] These alterations of both corneal epithelium and stroma might produce changes to a various extent of biomechanical properties of the cornea of oGVHD patients.

To date, few previous studies have investigated corneal biomechanics in patients with DED owing to different etiologies, but not oGVHD, by means of both ORA and CorVis ST.[24–27] The former is a non-contact tonometer that provides the measure of two parameters of corneal biomechanics: CRF that reflects the overall resistance of the cornea, and CH that measures the viscoelastic properties of the corneal tissue detecting the changes in the organization of collagen lamellae. CorVis ST is a device that employs an ultrahigh speed Scheimpflug camera to record dynamic deformation of the cornea. Impaired values of corneal biomechanics have been reported in DED patients with and without Sjögren’s syndrome[24–26]; conversely, Firat et al[27] reported no alterations of corneal biomechanical parameters in patients with DED. However, this study did not evaluate comprehensively ocular surface parameters, and did not stratify patients according to the severity of DED.

To the best of our knowledge, this is the first study evaluating corneal biomechanics in the setting of oGVHD. This condition is a type of iatrogenic DED, whose prevalence is increasing due to the widespread adoption of HSCT for the treatment of hematological disorders. We found significantly lower values of CH and CRF in oGVHD patients compared to healthy matched subjects. In addition, the severity of ocular surface impairment in these patients was associated with a greater alteration of corneal biomechanical properties, in particular for CH. This parameter reflects the viscoelastic properties of corneal tissue, detecting the changes in the organization of collagen lamellae. The most reasonable explanation to these findings is that the ocular surface inflammation caused by oGVHD might affect the corneal stroma ultrastructure by enhancing the rates of collagen and elastin enzymatic degradation. In particular, MMPs are proteolytic enzymes produced by stressed ocular surface epithelial cells, as well as by the immune cells that infiltrate these tissues. They are involved in the degradation of extracellular matrix components, and contribute to inflammation, wound healing, and tissue remodeling.[10–12] Overexpression of MMPs have been linked with corneal complications like ulcer and melting, that occur in the most severe stages of oGVHD, probably because of their role in the remodeling of extracellular matrix.[28–29]

Among MMP-family, MMP-9 represents the primary matrix-degrading enzyme of the ocular surface and is the only one that can be tested with a commercially available assay in clinical outpatient settings.[30] In the present study, we tested MMP-9 positivity in the tear film of patients with oGVHD and healthy subjects. All oGVHD patients resulted positive to MMP-9 assays against the negligible percentage of positivity among control subjects. This result is consistent with recent studies, which demonstrated increased tear levels of elastolytic enzymes, including MMP-9 and neutrophil elastase, in patients with oGVHD.[8,9] In addition, in order to overcome the drawback of the dichotomous result of MMP-9, which corresponds to either one of “ negative” or “ positive”, we employed a semi-quantitative method of grading the intensity of positivity, as already suggested by others.[31] Thus, we demonstrated that oGHVD patients exhibited different grades of MMP-9 positivity, and stronger positivity grade was associated with higher impairment of corneal biomechanics. This finding confirms the possible role of MMP-9 in the alteration of the stromal composition and organization also in the setting of oGVHD.

Conflicting results are available in the literature regarding central corneal thickness in DED. Although few studies showed similar values between DED patients and normal subjects[24–27], we found significantly lower values in patients with oGVHD compared to controls, in agreement with others.[32–34] The corneal thinning in DED is attributed to the increased osmolarity of the tear film that induces the activation of the inflammatory cascade and stimulate the production of high levels of the cytokine and MMP-9 by the distressed epithelial cells. These inflammatory events may lead to apoptotic death of surface epithelial cells of cornea and to destructive keratolysis and thinning.[35]

The main limitation of the study is related to the use of InflammaDry for the detection of pathological values of tear MMP-9. Although this is the only assay commercially available, a proteomic study with tear dosage of each type of MMPs would have provided a more accurate quantification of ocular surface inflammation, and its correlation with corneal biomechanics.

## Conclusion

The present study shows that oGVHD impairs biomechanical properties of the cornea, likely as a result of the alterations of ultrastructure and architecture of stromal collagen fibrils. Since corneal biomechanical alterations are closely related to the grade of ocular surface inflammation, they might represent a possible new surrogate marker of oGVHD severity.

## Founding

The authors received no specific funding for this work.

## Author contribution

G.G. conceptualization, project administration, writing – original draft. M.P. investigation, formal analisys. L.T. data curation, investigation. F.B. visualization, investigation. C.S. visualization, investigation. A.G. visualization, investigation E.C.C. validation, writing – review and editing. V.S. validation, writing – review and editing.

